# Undivided attention: The temporal effects of attention dissociated from decision, memory, and expectation

**DOI:** 10.1101/2021.05.24.445376

**Authors:** Denise Moerel, Tijl Grootswagers, Amanda K. Robinson, Sophia M. Shatek, Alexandra Woolgar, Thomas A. Carlson, Anina N. Rich

## Abstract

Selective attention prioritises relevant information amongst competing sensory input. Time-resolved electrophysiological studies have shown stronger representation of attended compared to unattended stimuli, which has been interpreted as an effect of attention on information coding. However, because attention is often manipulated by making only the attended stimulus a target to be remembered and/or responded to, many reported attention effects have been confounded with target-related processes such as visual short-term memory or decision-making. In addition, the effects of attention could be influenced by temporal expectation. The aim of this study was to investigate the dynamic effect of attention on visual processing using multivariate pattern analysis of electroencephalography (EEG) data, while 1) controlling for target-related confounds, and 2) directly investigating the influence of temporal expectation. Participants viewed rapid sequences of overlaid oriented grating pairs at fixation while detecting a “target” grating of a particular orientation. We manipulated attention, one grating was attended and the other ignored, and temporal expectation, with stimulus onset timing either predictable or not. We controlled for target-related processing confounds by only analysing non-target trials. Both attended and ignored gratings were initially coded equally in the pattern of responses across EEG sensors. An effect of attention, with preferential coding of the attended stimulus, emerged approximately 230ms after stimulus onset. This attention effect occurred even when controlling for target-related processing confounds, and regardless of stimulus onset predictability. These results provide insight into the effect of attention on the dynamic processing of competing visual information, presented at the same time and location.

## Introduction

To interact effectively in our environment, we need to select relevant information from the continuous stream of incoming visual information. It is important to understand how selective attention influences different stages of perceptual processing, to gain insight into the mechanisms by which the brain prioritises the important information. Time-resolved electrophysiological methods such as electroencephalography (EEG) and magnetoencephalography (MEG) have been used to study which stage of perceptual processing is affected by selective attention, using univariate analyses. For example, studies on spatial attention have found a stronger neural response to stimuli at a cued compared to an uncued location, starting around 80-100ms after stimulus onset (Eason, 1981; Eimer, 1994; Hillyard & Münte, 1984; Mangun & Hillyard, 1988; Neville & Lawson, 1987; Rugg et al., 1987). The time-course of feature-selective attention is generally thought to be slower compared to spatial attention, with a stronger neural response to the cued compared to the uncued feature around 100-150ms after stimulus onset (Czigler & Géczy, 1996; Eimer, 1997; Heslenfeld et al., 1997; Hillyard et al., 1998, but see Zhang & Luck (2009)).

A limitation of univariate methods is that they are unable to tease apart responses to cued and uncued information when it is presented at the same time and in the same location. In addition, univariate methods show only an overall increase or decrease in response, not how attention influences the information that is represented in that neural signal. In contrast, multivariate decoding methods can provide insight in the information that is coded in the brain, using the pattern of activation across sensors (MEG and EEG) or voxels (functional Magnetic Resonance Imaging) (e.g., Grootswagers et al., 2016; Haynes, 2015; Hebart & Baker, 2017). This method can be applied to MEG or EEG data to track the information coding of the cued and uncued stimuli over time, even when the stimuli are presented at the same time and location. Several studies have used multivariate decoding of time-resolved data to study the effects of attention on the coding of visual information (Battistoni et al., 2020; Goddard et al., 2019; Grootswagers et al., 2021; Kaiser et al., 2016; Moerel et al., 2021). These studies have found that the coding of cued information is sustained over time, whereas uncued information is represented only transiently and is not sustained over time. Although these previous studies have interpreted preferential coding of cued over uncued information as an effect of attention, their findings could be influenced by two other potential processes: target-related processes and expectation effects.

Target-related processes could differently influence the coding of cued and uncued information because attention to a stimulus is often manipulated by making the cued stimulus a target, while the uncued stimulus is not. This means that several other processes, which only occur for the cued stimulus, could drive the difference in coding of cued and uncued information. First, the cued target stimulus often has to be kept in visual short-term memory until the participant can make a decision. Second, the participant usually has to make decisions about the cued stimulus, but not about the uncued stimulus. For example, participants might have to categorise the shape of the cued object, whereas this is not the case for the uncued object (Goddard et al., 2019). Critically, the stimulus feature, such as object shape, that is kept in visual short-term memory and used to make a classification decision, is usually the same stimulus feature that is used in the decoding analysis. Thus, stronger information coding for cued compared to uncued stimuli, as found in previous work, might not be driven by attention alone. Either of these target-related processes could result in stronger information coding for the cued compared to uncued stimuli.

A few studies have tried to separate effects of attention from target-related processes by comparing the neural response to cued non-targets with uncued non-targets using either forward encoding models (Smout et al., 2019) or pattern classification (Grootswagers et al., 2021; Moerel et al., 2021). In the EEG study by Smout and colleagues (2019), participants viewed a sequence of oriented Gabor patches, while either searching for a target stimulus or monitoring the fixation dot for a colour change. Crucially, the authors did not analyse the neural response to targets, but rather to all non-target stimuli in the sequence, avoiding target-related confounds. The authors compared the coding of orientation information across two different sessions. Orientation information was task-relevant in a target search task performed in one session, but it was not task-relevant in a fixation change task performed in the other session. The results showed enhanced coding of orientation information in the target search task, compared to the fixation change task. However, the attentional breadth was likely larger for the target search task compared to the fixation change task. Grootswagers and colleagues (2021) presented the cued and uncued stimuli at the same time and location, while only analysing the neural response to non-targets. Participants monitored a sequence of letters overlaid on objects or *vice versa*, while they performed a 2-back target detection task on either the objects or the letters, comparing each item with the item that appeared two stimuli prior. The results showed enhanced of coding of the visual information that was currently relevant for the task (e.g., objects in the object condition) relative to irrelevant visual information. However, a 2-back task requires participants to hold the cued information in visual short-term memory. In addition, the cued and uncued stimuli were different sizes, which means the attentional breadth was different for the cued and uncued stimuli. To match the attentional breadth between the cued and uncued stimuli, both stimuli can be matched in size, and presented at the same location at the same time. In previous work, we used this method to compare the coding of the same orientations when they were either cued or uncued (Moerel et al., 2021). Importantly, the decision was an orthogonal dimension that was separated in time from the stimulus. The results showed enhanced coding of the cued compared to the uncued orientations, before participants could make a decision. However, participants had to keep the cued orientation in mind until the decision screen, which means that the results could be partly driven by visual short-term memory. Thus, it is still unclear whether effects of attention on visual processing occur when we control for the influence of all of these target-related processes.

In addition to target-related processes, the effects of selective attention on stimulus coding could be influenced by expectations about when a stimulus is likely to occur in time. Attention and temporal expectation could work together in an additive or interactive way to influence the processing of visual information (Kok et al., 2012; Smout et al., 2019; Zuanazzi & Noppeney, 2018). Temporal expectation can be experimentally manipulated through different temporal structures, such as through the use of explicit cues, or implicitly learned temporal rhythms (Nobre & Ede, 2018). Here we focus on implicitly learned rhythms of visual stimulus onset, as the explicit cues used to manipulate expectation can be very similar to those used in attention paradigms (see Rungratsameetaweemana et al., 2018 for a discussion). Some studies have found evidence of an effect of temporal expectation, as manipulated through rhythms, on the efficiency of early visual processing (Cravo et al., 2013; Rohenkohl et al., 2012). These studies found increased contrast sensitivity for visual stimuli presented at fixed compared to irregular intervals (Cravo et al., 2013; Rohenkohl et al., 2012), which was associated with increased phase entrainment of 1-4Hz oscillations. In addition, this entrainment was related to the behavioural discriminability of targets presented at a regular interval (Cravo et al., 2013). Although these studies do not investigate the interaction between temporal expectation and attention, they suggest that temporal expectation may interact with selective attention, as both processes affect visual processing.

Several studies have directly investigated the interaction between attention and expectation, but most of these studies focus on either spatial or stimulus feature expectations, that is, expectations about *where* or *what*, rather than *when* something will occur. These studies have found interactions between attention and expectation (Kok et al., 2012; Smout et al., 2019; Zuanazzi & Noppeney, 2018), as well as additive effects (Zuanazzi & Noppeney, 2018). Smout and colleagues (2019) investigated the interaction between attention and expectations about stimulus features. Although the authors found an interaction between attention and expectation on the coding of mismatch information, they did not find an interaction on the coding of visual orientation information. There is currently no consensus on whether temporal expectation is a possible confound in attention research, as little is known about the interaction between attention and implicitly learned temporal expectations. The lack of consensus is apparent from the variability in whether or not attention researchers make sure the stimulus onset is unpredictable to avoid temporal expectation confounds.

In this study, we investigated the time-course of the effect of attention on the coding of visual information in the brain. We compared the coding of attended and unattended visual stimuli that were presented simultaneously in time and space, while 1) making sure the pattern classification could not be driven by target-related processing, and 2) directly investigating the influence of temporal expectation. We recorded EEG data while participants performed a target detection task on sequences of central stimuli comprised of overlaid oriented blue and orange lines. Each sequence was cued by a stimulus of a specific colour and orientation. Participants had to respond when this cued target appeared in the sequence. This task ensured that one colour was cued (attended) and one colour was uncued (unattended) for each sequence. We used multivariate pattern analysis (MVPA) to compare the orientation coding of the same stimuli when they were attended or not. Importantly, we controlled for effects of visual short-term memory and decision-making by ensuring these effects could not drive classification performance. Specifically, we minimised effects of visual short-term memory by using a target detection task (effectively a “0-back” task), so participants did not have to retain the decoded stimulus orientation over a delay period. Participants did have to remember which was the target orientation (horizontal or vertical) for the duration of the sequence, but this could not drive the classification, as the target orientation was not informative about the decoded stimulus orientation. That is, target orientations were never the same as the decoded stimulus orientations, and all decoded stimulus orientations were presented an equal number of times when participants were looking for a horizontal compared to a vertical target. Critically, we only analysed the non-targets. Likewise, decision-making processes could not drive the classifier. Although a decision had to be made on each trial, it was always the same decision about whether the stimulus was a target. Because all decoded stimulus orientations were presented an equal number of times when participants were looking for a horizontal compared to a vertical target, the decisions were matched between all decoded stimulus orientations. To examine the influence of temporal expectation on attention effects, we asked 1) whether there was an effect of temporal predictability on the coding of visual information, and 2) whether an effect of attention on the coding of visual information occurred regardless of whether the stimulus onset was temporally predictable or not. To this end, we compared the orientation coding of stimuli presented at a constant (predictable) stimulus onset (interstimulus interval (ISI) = 200ms) with that from varied (unpredictable) stimulus onset (ISI between 100ms and 300ms). Together, we used these manipulations to examine the effect of attention on stimulus coding, while controlling for visual short-term memory and decision-making confounds, and to explore the effect of temporal expectancy. Our results showed preferential coding of the cued compared to the uncued information emerging from about 230ms after stimulus onset. This difference is likely driven by the prioritisation of the task-relevant information through attention, as our manipulations made sure that visual short-term memory or decision-making could not drive the classification, and it did not interact with temporal predictability.

## Methods

### Participants

Twenty healthy adults participated in this study (12 female/8 male; 19 right-handed/1 left-handed; mean age = 23.80 years; age range = 18-59 years). All participants reported normal or corrected-to-normal vision and normal colour vision. Participants were recruited from the University of Sydney and received $40 AUD for their participation. The study was approved by the University of Sydney ethics committee and informed consent was obtained from all participants.

### Stimuli and design

The stimuli consisted of approximately equiluminant blue (RGB = 239, 159, 115) and orange (RGB = 72, 179, 217) oriented lines, overlaid at fixation, presented on a mid-grey background (Figure 1A). The oriented lines were phase randomised over trials, had a spatial frequency of 1.15 cycles/degree, and were shown within a circular aperture with a diameter of 5.20 degrees of visual angle. There were 4 possible line orientations for the non-target stimuli, which were used for analysis: 22.5°, 67.5° 112.5°, and 157.5°. Orange and blue lines were always shown together, rotated 45° with respect to each other, resulting in 8 unique combinations of orientations (Figure 1B). These orientations were chosen to make sure they were orthogonal over attention conditions for the decoding analysis.

**Figure 1.**
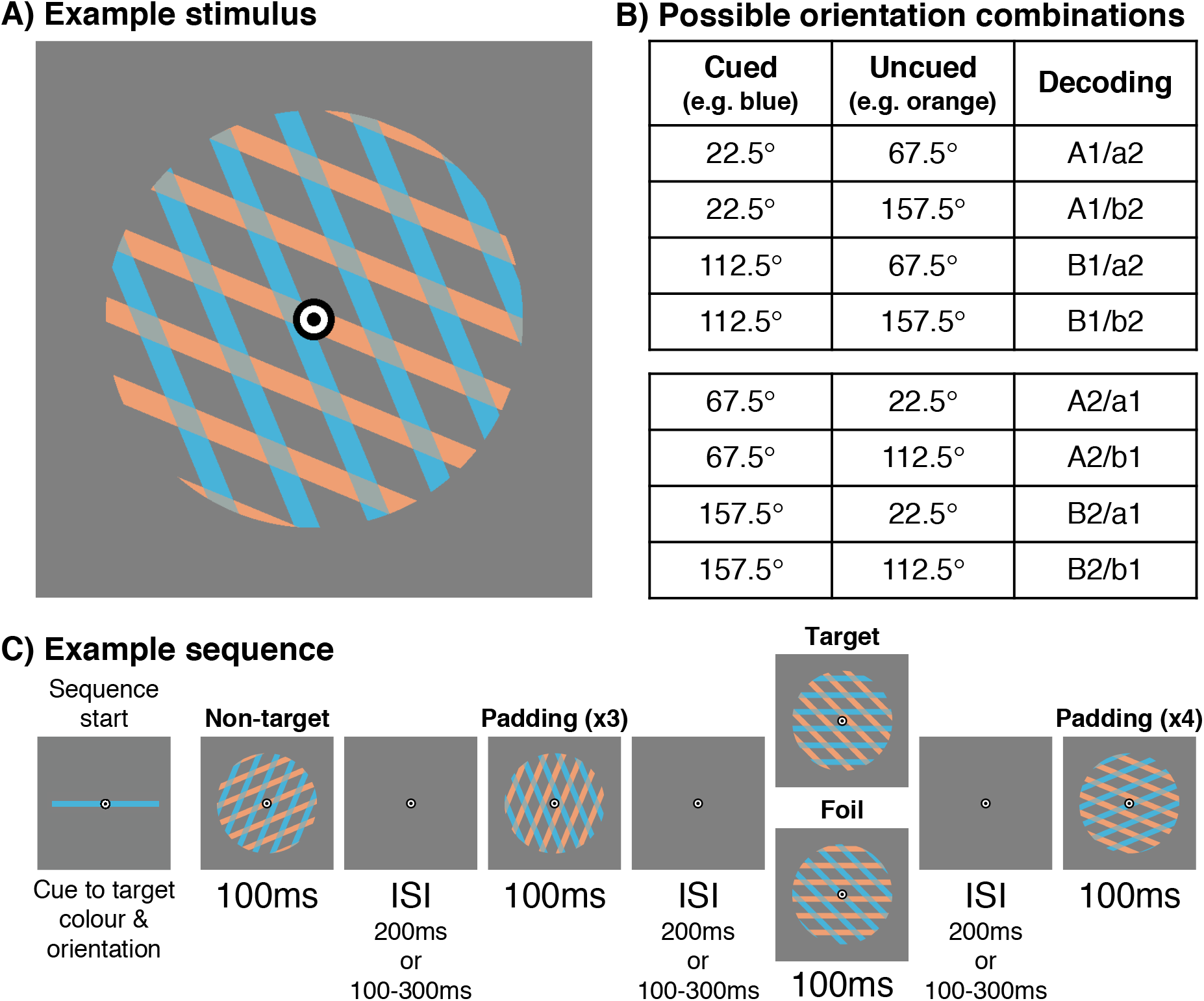
Experimental design. A) The stimuli consisted of blue and orange oriented lines, overlaid at fixation. B) There were 4 possible line orientations: 22.5°, 67.5° 112.5°, 157.5°. The blue and orange lines were always rotated 45°, resulting in 8 possible combinations of orientations. We made sure the cued and uncued orientations were orthogonal in the decoding analysis by dividing the orientations into 2 pairs, labelled 1 and 2 in the “Decoding” column. The “Decoding” column in the table shows a label for cued (capital letter) and uncued orientation (lower case letter). For the cued orientation, we decoded A vs. B for each pair (A1 vs. B1 and A2 vs. B2), and for the uncued orientation, we decoded a vs. b for each pair (a1 vs. b1 and a2 vs. b2). Note that for each cued orientation, both uncued orientations occur equally often and vice versa. For example, the cued orientation A1 is equally often paired with a2 as b2. C) Shows an example of part of a sequence. At the start of each sequence, participants saw the target colour and orientation for that sequence until they pressed a key to start. Stimuli were presented for 100ms. Each stimulus consisted of lines of the cued and uncued colour. The ISI for the sequence was either constant at 200ms or varied between 100ms and 300ms. The third stimulus in the sequence shows an example of a target at the top (cued blue target orientation), and a foil at the bottom (uncued orange target orientation). Participants had to respond to targets but not to foils. The target and foil trials, along with 3 padding trials before and 4 padding trials after each target or foil, were not used in the analysis. Only the non-target stimuli (the first stimulus in this example) were used in the analysis.

Participants were instructed to maintain fixation on a central bullseye throughout the entire experiment and to respond to a target presented at fixation using a button press response. The stimuli were presented in 64 sequences. At the start of each sequence, participants were shown the target stimulus. The target had a specific orientation (0° or 90°) and colour (orange or blue) and was counterbalanced over sequences. Participants were instructed to attended to the lines in the target colour and ignore the lines in the other colour. The task was to press a button as soon as lines in the target orientation and colour appeared. The purpose of the task was to keep participants engaged, and to make sure participants attended to the lines of one colour, ignoring the other colour. In each sequence, the target orientation (0° or 90°) was paired with either 45° or 135° in the other colour. The target orientation could either be shown in the cued colour (target event), or in the uncued colour (foil event), and participants had to respond only to the target events. For instance, if the target was 90°-blue, the possible target stimuli for that sequence were 90°-blue/45°-orange, and 90°-blue/135°-orange, while the possible foils for that sequence were 45°-blue/90°-orange, and 135°-blue/90°-orange (see Figure 1C).

Each sequence consisted of 104 non-target stimulus presentations. In addition, there were 1 or 2 target events and 1 or 2 foil events per sequence, counterbalanced over sequences. In addition, non-target padding trials were added at the start and end of each sequence as well as around targets and foils. These padding trials were randomly drawn without replacement from the 8 possible trial types (Figure 1B) and were removed from further analysis. There were 4 padding trials at the start and end of each sequence, 3 padding trials before each target and foil, and 4 after. The total number of stimulus presentations in each sequence ranged between 128 and 144, depending on the number of targets and foils in each sequence. All target events, foil events, and padding were removed, and only the 104 non-target stimulus presentations in each sequence were used for further analysis.

Each stimulus in the sequence was presented for 100ms, while the interstimulus interval (ISI) duration was either 200ms in the constant ISI condition or varied between 100ms and 300ms in the varied ISI condition (Figure 1C). In the varied ISI condition, there were 13 possible ISIs, equally spaced between 100ms and 300ms, reflecting the 60Hz refresh rate of the monitor (i.e., 16.67ms steps). We counterbalanced over sequences whether the ISI was constant or varied. Within a sequence, the 104 stimulus presentations were balanced for every combination of ISI before stimulus onset (13 for variable ISI condition) x orientation (4) x attention condition (2).

Participants completed 3 training parts during the EEG setup. Training parts 1 and 2 consisted of 4 sequences with 32 stimuli per sequence. In the first training part, participants only saw lines of the cued colour and the task was slowed down to 400ms stimulus presentation and 400ms ISI. In the second training part, the lines in the uncued colour were introduced, while the timing was kept the same as part 1. The third training part consisted of 8 sequences with 104 stimulus presentations per sequence. This training part was the same as the experiment: the stimulus presentation was sped up to 100ms, and the ISI was either 200ms (4 sequences) or varied from 100 to 300ms (4 sequences).

### EEG acquisition and pre-processing

We recorded continuous EEG data from 64 electrodes, digitised at a sample rate of 1000-Hz, using a BrainVision ActiChamp system. The electrodes were arranged according to the international standard 10–10 system for electrode placement (Oostenveld & Praamstra, 2001) and were referenced online to Cz. We performed offline pre-processing using the EEGlab toolbox in Matlab (Delorme & Makeig, 2004), using the same pre-processing pipeline as in earlier decoding work (Grootswagers et al., 2021; Robinson et al., 2020). We filtered the data using a Hamming windowed FIR filter with 0.1Hz high pass and 100Hz low pass filters, re-referenced the data to an average reference, and down-sampled the data to 250Hz. We created epochs from −100ms to 800ms relative to stimulus onset.

### Decoding analysis

We conducted 3 different decoding analyses, to test for 1) an effect of attention on orientation coding, 2) an effect of temporal expectation on orientation coding, and 3) an interaction between attention and temporal expectation. To investigate the time-course of the effect of attention on the coding of visual information in the brain, we collapsed the data over the different timing predictability conditions. We used MVPA to determine whether the pattern of activation across EEG channels carried information about the cued orientation and the uncued orientation for each time-point. Comparing the coding of the cued and uncued orientation allowed us to determine at which time-points an effect of attention occurred, as indicated by stronger coding of cued compared to uncued orientations. We used the CoSMoMVPA toolbox for MATLAB for all decoding analyses (Oosterhof et al., 2016). For decoding of the cued orientation, we trained a regularised Linear Discriminant Analysis classifier to distinguish between orientations for each time-point in the epoch. Because the cued and uncued orientations were presented simultaneously, we made sure orientations were orthogonal over attention conditions. We did this by dividing the 4 possible cued orientations into 2 pairs, 22.5° vs. 112.5° and 67.5° vs. 157.5°, and running the decoding analysis within a single pair. For instance, when the cued orientations were 22.5° and 112.5°, the uncued orientations could be either 67.5° or 157.5°, each occurring equally often for each cued orientation (Figure 1B). We averaged decoding accuracies across pairs, resulting in a chance level of 50%. We determined decoding performance for each individual participant using a leave-one-sequence-out cross-validation approach, and then analysed the subject-averaged results at the group level. The decoding analysis of the uncued orientation was conducted in the same way. For plotting purposes, we bootstrapped 95% confidence intervals across participants using 10,000 bootstrap samples.

To examine the effect of temporal expectation, we split the data into the constant and variable ISI conditions, and performed the analysis described above separately for both the attention (cued or uncued) and temporal expectation (constant ISI or varied ISI) conditions, and then averaged across the cued and uncued conditions to obtain a single time-course of decoding accuracies per temporal predictability condition.

To investigate whether the effect of attention was 1) present, and 2) the same, for the two ISI conditions, we examined the orientation decoding accuracies separately for the different attention (cued or uncued) x temporal expectation (constant ISI or varied ISI) conditions.

### Statistical inference

We used Bayesian statistics to determine the evidence for above-chance decoding (alternative hypothesis) and chance decoding (null hypothesis) for each point in time (Dienes, 2011; Kass & Raftery, 1995; Morey et al., 2016; Rouder et al., 2009), using the Bayes Factor R package (Morey & Rouder, 2018). We did this for 1) the orientation coding separated for the cued and uncued conditions, 2) the orientation coding separated for the constant and varied ISI conditions, and 3) the orientation coding separated for both the attention and ISI conditions. We used a half-Cauchy prior for the alternative hypothesis, centred around chance (d = 0, i.e., 50% decoding accuracy), with the default width of 0.707 (Jeffreys, 1998; Rouder et al., 2009; Wetzels et al., 2011) to capture directional (above chance) effects. We excluded the interval ranging from d = 0 to d = 0.5 from the prior to exclude irrelevant effect sizes, and used a point null at d =0.

We tested for an effect of attention for data combined across temporal predictability conditions, and separately for the constant and varied timing conditions, by calculating the difference between decoding accuracies for the cued and uncued orientations. Because we had the *a-priori* hypothesis that the orientation coding of the cued orientation would be higher compared to uncued orientation, we used the same half-Cauchy (directional) prior described above, centred around 0, to test whether decoding accuracies of the cued orientation were higher than for the uncued orientation. We tested for an effect of temporal expectation by comparing the orientation coding of stimuli presented in the constant and varied ISI conditions, collapsed over attention conditions. We used the half-Cauchy prior described above to test whether orientation decoding accuracies in the constant ISI condition were higher than those in the varied ISI condition. To assess whether the effect of attention was different across the two ISI conditions, we compared the difference between decoding accuracies for the cued and uncued orientations directly for the constant and varied ISI conditions. Unlike the tests described above, we did not have an *a-priori* prediction about the direction of this effect. Therefore, we used whole Cauchy (not directional) prior for the alternative, and excluded an interval ranging from effect size of −0.5 to 0.5 from the prior to make it comparable to the other analyses.

Bayes factors (BF) above 1 indicate evidence in the direction of the alternative hypothesis, and Bayes factors below 1 indicate evidence in the direction of the null hypothesis. Bayes factors below 1/3 or above 3 are usually interpreted as substantial evidence, and Bayes factors below 1/10 or above 10 are usually interpreted as strong evidence (Wetzels et al., 2011). We defined the onset time of above chance decoding as the second consecutive time-point with a Bayes factor above 10 (strong evidence).

## Results

The purpose of the task was to make sure participants attended to the orientations in the cued colour, while ignoring the orientations in the other colour. The task was difficult due to its fast nature, ensuring participants had to pay attention. Responses within 2000ms after the onset of the target or foil stimulus were counted as hits or false alarms respectively. Participants performed the target-detection task in the constant and varied interval conditions respectively with a mean hit rate of 60.96% and 58.06% (SD = 11.60% and 11.49%), a mean false alarm rate on foils (target orientation in uncued colour) of 11.12% and 9.89% (SD = 6.46% and 6.43), and a mean false alarm rate on the non-target trials of 0.34% and 0.34% (SD = 0.41% and 0.45%). This indicates that the participants were engaged in the task and managed to discriminate between the orientations of the cued and uncued colour.

We decoded the cued and uncued orientations for each time-point to determine whether there is an effect of attention on the coding of visual information (Figure 2A). There was strong evidence for above chance decoding of both the cued and uncued orientation from approximately 80ms after stimulus onset (84ms and 80ms respectively). There was strong evidence for an effect of attention, defined as stronger coding of the cued compared to uncued orientation, from 232ms (Figure 2B). This suggests the brain initially represented both the cued and uncued stimuli to a similar extent, but later prioritised the processing of the attended information.

**Figure 2.**
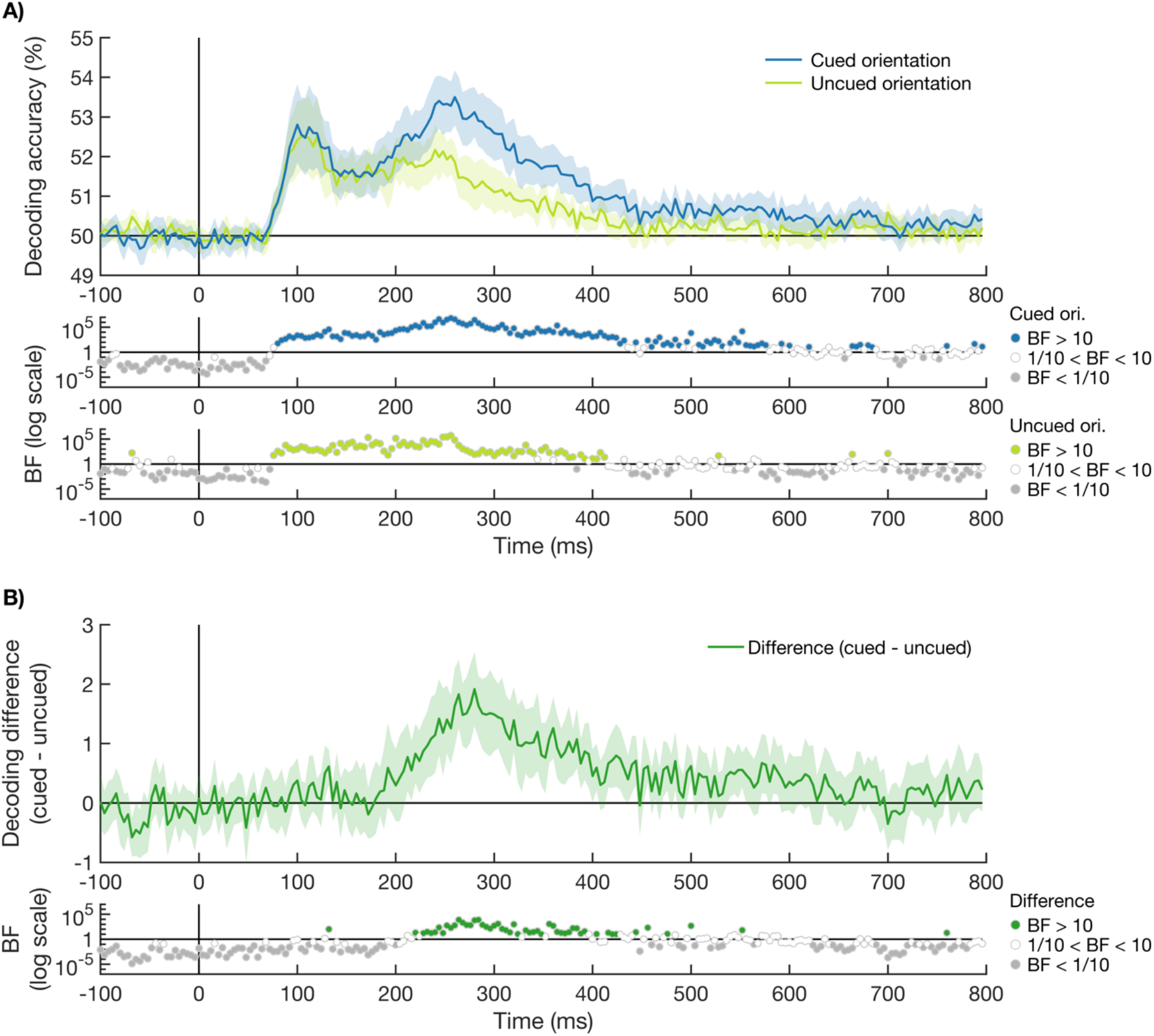
Time-course of decoding accuracies of the cued and uncued orientations. A) shows the decoding accuracy of the stimulus orientation over time, when the orientation was either presented in the cued colour (blue) or uncued (light green). The stimulus onset is marked with a vertical black line at 0ms. Theoretical chance is 50% decoding accuracy, and shaded areas around the plot lines show the bootstrapped 95% confidence interval across participants (N = 20). Bayes factors are given below the plot on a log scale. Bayes factors below 1/10 are shown in grey, indicating strong evidence for the null hypothesis. Bayes factors above 10 are shown in the plot colour (blue for cued and light green for uncued), indicating strong evidence for the alternative hypothesis. Bayes factors between 1/10 and 10 are shown in white. The cued orientation could be decoded from 84ms after the onset of the stimulus, and the uncued orientation from 80ms after stimulus onset. B) shows the effect of attention, measured as the difference in orientation decoding between the cued and uncued orientation (cued - uncued). The Bayes factors are given below the plot on a log scale. An effect of attention was present from 232ms after stimulus onset.

The second aim of this study was to examine the influence of temporal expectation on the effect of attention. First, we determined whether there was a main effect of temporal predictability on the processing of visual orientation information, and then we tested whether there was an interaction between attention and temporal predictability. To determine whether there was a main effect of temporal expectation, we compared the decoding of orientation between stimuli presented in the constant and varied ISI sequences, collapsing across attention conditions. The decoding accuracies for the constant and varied ISI conditions are shown in Figure 3A. We calculated the effect of temporal expectation as the difference between orientation coding for stimuli in the constant and varied ISI sequences (see Figure 3B). There was no reliable effect of temporal expectation on the processing of visual information. Although a few time-points after 300ms showed evidence for stronger orientation coding for the constant compared to varied ISI conditions (BF_10_ > 10), most time-points after 300ms showed either inconclusive evidence or strong evidence for no effect (BF_10_ < 1/10).

**Figure 3.**
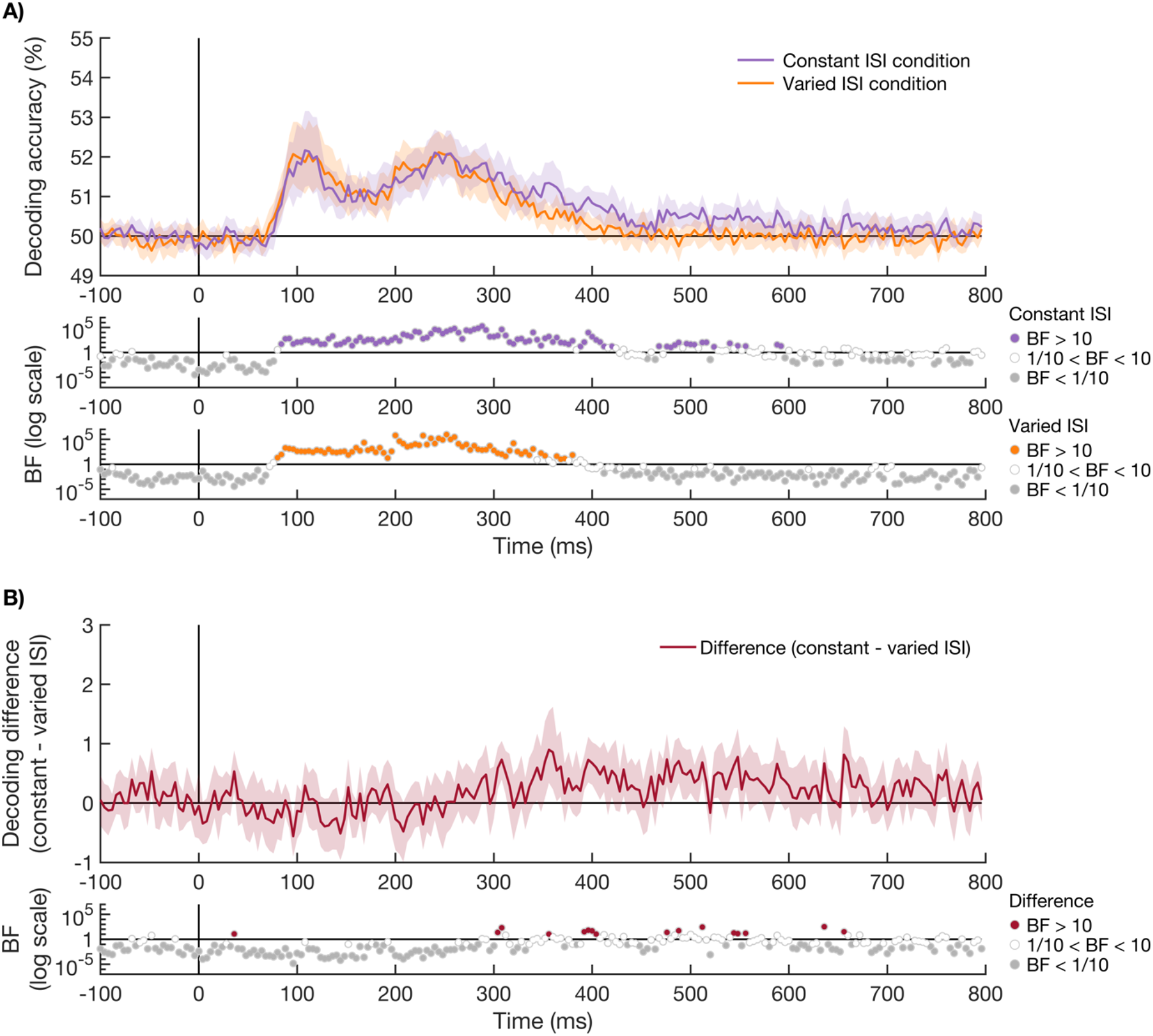
Time-course of decoding accuracies of the orientations presented at a constant and varied ISI. A) shows orientation decoding accuracy of stimuli presented at a constant ISI (purple) or varied ISI (orange). Plotting conventions are the same as in Figure 2. The orientations of stimuli presented at a constant ISI or varied ISI could be decoded from 88ms and 84ms after stimulus onset respectively. B) shows the effect of temporal expectation, measured as the difference in orientation decoding between the stimuli presented at a constant or varied ISI (constant - varied ISI). The BFs are given below the plot on a log scale. There was no effect of temporal expectation on the coding of orientation information. A few time-points after 300ms showed stronger orientation coding for the constant compared to varied ISI conditions, but the Bayes factors for most time-points in the 300ms to 800ms time-window either suggested inconclusive evidence or evidence for no difference.

To examine whether an effect of attention was present regardless of timing predictability, we compared the decoding of the cued and uncued orientations separately for the constant ISI sequences (Figure 4A) and the varied ISI sequences (Figure 4B). We observed an effect of attention for both timing conditions, starting at 248ms after stimulus onset for the constant timing condition, and at 232ms after stimulus onset for the varied timing condition. We directly compared the effects of attention in the two different timing conditions, to examine the interaction between attention and temporal expectation (Figure 4C). Although a few time-points showed evidence for a larger effect of attention in the varied ISI condition, no two consecutive time-points showed strong evidence (BF_10_ > 10) for this difference, and the majority of time-points showed strong evidence for no effect (BF_10_ < 1/10). Note that although the visual input was perfectly matched between the constant and varied ISI conditions until 200ms after stimulus onset, this is not the case after this time. While the next stimulus in the sequence was always presented 300ms after the preceding one in the constant ISI condition, it was presented between 200ms and 400ms after stimulus onset in the varied ISI condition (see grey line in figure 4A and grey shaded area in figure 4B). This means that masking by the next stimulus in the sequence occurred earlier in some of the varied ISI trials (and later in others) compared to the constant ISI trials.

**Figure 4.**
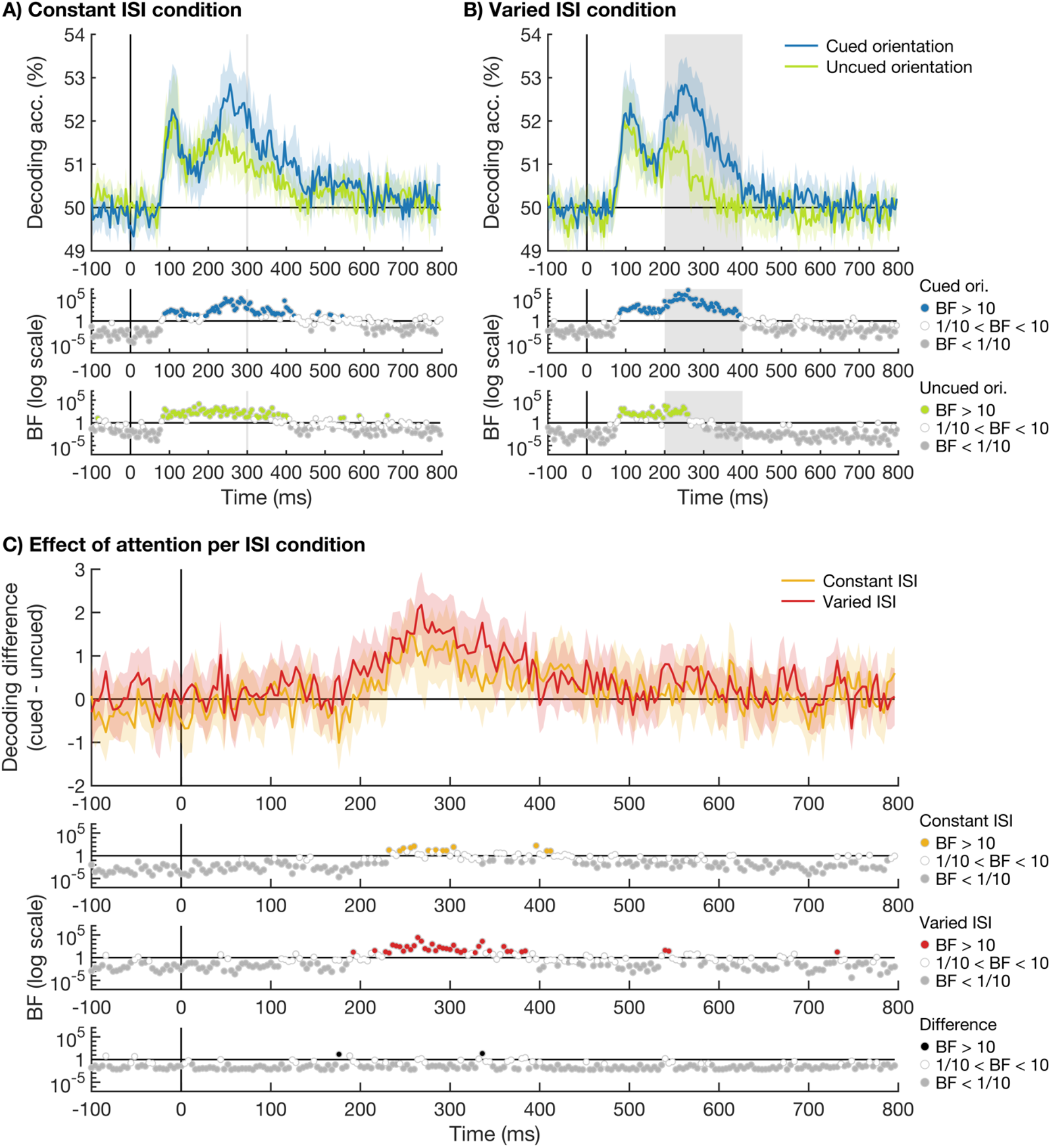
Time-course of decoding accuracies of the cued and uncued orientations for the varied and constant ISI conditions. Plots show the decoding accuracies of the cued (blue) and uncued (light green) stimulus orientation over time for the constant ISI condition in A) and the varied ISI condition in B). The possible onset time(s) of the next stimulus in the sequence are indicated by a grey shaded line at 300ms in A) and a grey shaded area between 200ms and 400ms in B). C) shows the effect of attention on stimulus coding, calculated as the difference in decoding accuracy for the cued and uncued conditions, for the constant ISI condition (yellow) and the varied ISI condition (red). We observed an effect of attention for both timing conditions, starting at 248ms after stimulus onset for the constant ISI condition, and at 232ms after stimulus onset for the varied ISI condition. We directly compared the effect of attention for the 2 different ISI conditions, and found no 2 consecutive time-points with BF_10_>10. Plotting conventions are the same as in Figure 2.

## Discussion

In this experiment we investigated the time-course of the effect of attention on visual information processing. We controlled for effects of target related processes, such as visual short-term memory and decision-making, by using a simple target detection task and focusing the analysis on the non-target items in the sequence. In addition, we investigated whether temporal expectation modulated the effects of attention on information processing by presenting the stimuli with either a constant (predictable) interval between stimuli of 200ms, or a varied (unpredictable) interval between 100ms and 300ms. The results showed that whereas the cued and uncued visual input was initially coded equally from about 80-84ms after stimulus onset, an effect of attention emerged from approximately 232ms. After this time, the cued visual information was more strongly coded than uncued information. We observed this effect of attention, with a similar time-course, in both the varied and constant ISI conditions. These findings show that attention selectively enhances the relevant compared to irrelevant visual information, which is consistent with the role of attention in boosting and maintaining task-relevant information. Importantly, this effect of attention on stimulus coding occurred even when we controlled for the effects of visual short-term memory and decision-making, and occurred regardless of whether the onset timing was predictable or not.

Our findings of an effect of attention on orientation coding are consistent with previous studies that used time-resolved methods in combination with MVPA. These studies found more sustained coding for the cued compared to uncued features and/or objects (Goddard et al., 2019; Grootswagers et al., 2021; Moerel et al., 2021). In line with the study by Moerel and colleagues (2021), we show that these effects are not contingent on confounding stimulus and decision. The current study builds on previous findings by showing that attention effects occur even when we control for target-related processing and ensure that decisions do not confound the stimulus classification, and that they occur regardless of whether the stimulus onset timing is predictable or not. One difference with previous research is how long the cued information was maintained in the neural signal, as this appears to be sustained for a shorter period in this study compared to previous findings (Goddard et al., 2019; Grootswagers et al., 2021; Moerel et al., 2021). This difference could be caused by a number of different processes. First, the masking of each stimulus by the subsequent stimulus occurs earlier in this paradigm compared to previous experiments (Goddard et al., 2019; Grootswagers et al., 2021; Moerel et al., 2021; Smout et al., 2019), and masking has been shown to affect the coding of visual information (Robinson et al., 2019). Secondly, this difference could be driven by the reduced requirement to maintain the cued visual information over time compared to some of the previous paradigms (Grootswagers et al., 2021; Moerel et al., 2021). Third, compared to other studies, there is a reduced requirement to make complex decisions about the cued information (Goddard et al., 2019; Moerel et al., 2021), which could affect the decoding duration. These different explanations are not mutually exclusive, and all seem likely to play a role. Although the decoding duration of cued information might be dependent on the study design, the effect of attention emerges at a similar time across studies, approximately 200-300ms after stimulus onset. The similar timing of the emergence of preferential processing of the relevant compared to irrelevant information is in line with this effect being driven by attention. Our study adds to previous findings by unequivocally demonstrating the temporal dynamics of attention on the processing of visual information, without target-related processing confounds. In addition, our design allows us to study attended and unattended stimuli presented simultaneously in the same spatial location, in a fast and efficient paradigm.

We found evidence that temporal predictability did not interact with the effect of attention on visual processing for the timescale used in this paradigm: a constant ISI of 200ms and a varied ISI of 100-300ms. The stimulus presentation was kept constant at 100ms, resulting in a stimulus onset asynchrony of 300ms, and 200-400ms respectively. This finding fits with findings from Smout and colleagues (2019), who showed that expectation about stimulus identity did not interact with the effect of attention on stimulus coding. Our study adds to these findings by investigating the contribution of temporal expectation. In addition, we did not find a main effect of temporal expectation on orientation processing. Although a few time-points after 300ms showed evidence for stronger orientation coding in the constant compared to varied ISI condition, likely driven by more sustained decoding of the uncued orientation in the constant condition, most time-points showed either inconclusive evidence or evidence for no difference. The difference in those few time-points is likely attributable to the difference in visual stimulation starting after 200ms. The next stimulus, which is likely to cause masking, occurred earlier in some of the varied ISI trials compared to the constant ISI trials. The lack of a main effect of temporal expectation does not seem in agreement with other studies that found evidence for enhanced visual processing for stimuli presented at a constant compared to varied ISI (Cravo et al., 2013; Rohenkohl et al., 2012). However, our results are not directly comparable to these previous studies as we used a different method to assess the effect of temporal expectation on the processing of visual information. Whereas we used MVPA of EEG data to track the coding of visual information, previous studies have investigated the effect of implicit temporal expectations on phase-locking of oscillations (Cravo et al., 2013) and behaviour (Cravo et al., 2013; Rohenkohl et al., 2012). Our methods are more comparable to those used by Smout and colleagues (2019), who showed that expectation about stimulus identity did not affect orientation coding.

One possible explanation for the apparent difference between our results and other studies of the effects of temporal expectation (Cravo et al., 2013; Rohenkohl et al., 2012) could be the timescale, as other studies have used a slower ISI in the predictable timing condition (400ms compared to 200ms), as well as a wider range of range of possible ISIs for the unpredictable timing condition (200-600ms instead of 100-300ms) (Cravo et al., 2013; Rohenkohl et al., 2012). It is possible that this, perhaps stronger, manipulation of temporal expectation would lead to different results in our paradigm, as this would give participants more time to process the current stimulus and prepare for the next stimulus. Another, not mutually exclusive, explanation is that the perceptual difficulty of the stimuli plays a role in whether temporal attention can affect the processing of visual information. Although our task was difficult due to its fast nature, the *perceptual* difficulty was low, as the non-targets were always rotated at least 22.5° from the target orientation and there was no added visual noise. This was not the case for some of the previous studies, where target orientations were embedded in visual noise (Cravo et al., 2013; Rohenkohl et al., 2012). It is possible that the effect of temporal expectation may have a larger effect under conditions of high perceptual difficulty, when there is more to gain from attending precisely in time. Future work could disentangle whether temporal expectation effects are dependent on perceptual difficulty and longer time scales. In addition, future work could also compare a predictable timing condition to a condition where temporal expectation is violated, rather than a condition where there is no temporal expectation as in our varied condition. It is possible that an interaction would occur when temporal expectations are violated, in line with the findings of Smout and colleagues (2019), who found an interaction between attention and expectation about stimulus features on the processing of mismatch information only.

Taken together, our results showed that attention selectively enhanced the high-level processing of stimulus relevant features from about 232ms after stimulus onset, without target-related processing confounds. The selection of task-relevant information occurred even when the stimulus presentation was fast, and occurred regardless of whether the onset was temporally predictable or not. These findings reveal a detailed picture for the time-course of the prioritisation of task-relevant information, in the presence of competing information at the same time and location.

## Acknowledgements

This work was supported by an International Research Training Program Scholarships from Macquarie University awarded to DM, an Australian Research Council (ARC) Discovery Early Career Researcher Award awarded to AKR (DE200101159), ARC Discovery Projects awarded to TAC (DP160101300 and DP200101787) and to ANR and AW (DP170101840), and by the Medical Research Council (UK) intramural funding (SUAG/052/G101400) awarded to AW.

## References

Battistoni, E., Kaiser, D., Hickey, C., & Peelen, M. V. (2020). The time course of spatial attention during naturalistic visual search. Cortex, 122, 225–234. https://doi.org/1fi0.1016/j.cortex.2018.11.018

Cravo, A. M., Rohenkohl, G., Wyart, V., & Nobre, A. C. (2013). Temporal Expectation Enhances Contrast Sensitivity by Phase Entrainment of Low-Frequency Oscillations in Visual Cortex. Journal of Neuroscience, 33(9), 4002–4010. https://doi.org/10.1523/JNEUROSCI.4675-12.2013

Czigler, I., & Géczy, I. (1996). Event-related potential correlates of color selection and lexical decision: Hierarchical processing or late selection? International Journal of Psychophysiology, 22(1–2), 67–84. https://doi.org/10.1016/0167-8760(96)00005-0

Delorme, A., & Makeig, S. (2004). EEGLAB: An open source toolbox for analysis of single-trial EEG dynamics including independent component analysis. Journal of Neuroscience Methods, 134(1), 9–21. https://doi.org/10.1016/j.jneumeth.2003.10.009

Dienes, Z. (2011). Bayesian Versus Orthodox Statistics: Which Side Are You On? Perspectives on Psychological Science, 6(3), 274–290. https://doi.org/10.1177/1745691611406920

Eason, R. G. (1981). Visual evoked potential correlates of early neural filtering during selective attention. Bulletin of the Psychonomic Society, 18(4), 203–206. https://doi.org/10.3758/BF03333604

Eimer, M. (1994). “Sensory gating” as a mechanism for visuospatial orienting: Electrophysiological evidence from trial-by-trial cuing experiments. Perception & Psychophysics, 55(6), 667–675. https://doi.org/10.3758/BF03211681

Eimer, M. (1997). An event-related potential (ERP) study of transient and sustained visual attention to color and form. Biological Psychology, 44(3), 143–160. https://doi.org/10.1016/S0301-0511(96)05217-9

Goddard, E., Carlson, T. A., & Woolgar, A. (2019). Spatial and feature-selective attention have distinct effects on population-level tuning. BioRxiv, 530352. https://doi.org/10.1101/530352

Grootswagers, T., Robinson, A. K., Shatek, S. M., & Carlson, T. A. (2021). The neural dynamics underlying prioritisation of task-relevant information. Neurons, Behavior, Data Analysis, and Theory, 2020.06.25.172643. https://doi.org/10.51628/001c.21174

Grootswagers, T., Wardle, S. G., & Carlson, T. A. (2016). Decoding Dynamic Brain Patterns from Evoked Responses: A Tutorial on Multivariate Pattern Analysis Applied to Time Series Neuroimaging Data. Journal of Cognitive Neuroscience, 29(4), 677–697. https://doi.org/10.1162/jocn_a_01068

Haynes, J.-D. (2015). A Primer on Pattern-Based Approaches to fMRI: Principles, Pitfalls, and Perspectives. Neuron, 87(2), 257–270. https://doi.org/10.1016/j.neuron.2015.05.025

Hebart, M. N., & Baker, C. I. (2017). Deconstructing multivariate decoding for the study of brain function. NeuroImage. https://doi.org/10.1016/j.neuroimage.2017.08.005

Heslenfeld, D. J., Kenemans, J. L., Kok, A., & Molenaar, P. C. M. (1997). Feature processing and attention in the human visual system: An overview. Biological Psychology, 45(1), 183–215. https://doi.org/10.1016/S0301-0511(96)05228-3

Hillyard, S. A., & Münte, T. F. (1984). Selective attention to color and location: An analysis with event-related brain potentials. Perception & Psychophysics, 36(2), 185–198. https://doi.org/10.3758/BF03202679

Hillyard, S. A., Teder-Sälejärvi, W. A., & Münte, T. F. (1998). Temporal dynamics of early perceptual processing. Current Opinion in Neurobiology, 8(2), 202–210. https://doi.org/10.1016/S0959-4388(98)80141-4

Jeffreys, H. (1998). The Theory of Probability. OUP Oxford.

Kaiser, D., Oosterhof, N., & Peelen, M. (2016). The Neural Dynamics of Attentional Selection in Natural Scenes. The Journal of Neuroscience : The Official Journal of the Society for Neuroscience, 36, 10522–10528. https://doi.org/10.1523/JNEUROSCI.1385-16.2016

Kass, R. E., & Raftery, A. E. (1995). Bayes Factors. Journal of the American Statistical Association, 90(430), 773–795. https://doi.org/10.1080/01621459.1995.10476572

Kok, P., Rahnev, D., Jehee, J. F. M, Lau, H. C., Lange, D., & P, F. (2012). Attention Reverses the Effect of Prediction in Silencing Sensory Signals. Cerebral Cortex, 22(9), 2197–2206. https://doi.org/10.1093/cercor/bhr310

Mangun, G. R., & Hillyard, S. A. (1988). Spatial gradients of visual attention: Behavioral and electrophysiological evidence. Electroencephalography and Clinical Neurophysiology, 70(5), 417–428. https://doi.org/10.1016/0013-4694(88)90019-3

Moerel, D., Rich, A. N., & Woolgar, A. (2021). Selective attention and decision-making have separable neural bases in space and time. BioRxiv, 2021.02.28.433294. https://doi.org/10.1101/2021.02.28.433294

Morey, R. D., Romeijn, J.-W., & Rouder, J. N. (2016). The philosophy of Bayes factors and the quantification of statistical evidence. Journal of Mathematical Psychology, 72, 6–18. https://doi.org/10.1016/j.jmp.2015.11.001

Morey, R. D., & Rouder, J. N. (2018). BayesFactor: Computation of Bayes Factors for Common Designs. https://CRAN.R-project.org/package=BayesFactor

Neville, H. J., & Lawson, D. (1987). Attention to central and peripheral visual space in a movement detection task: An event-related potential and behavioral study. I. Normal hearing adults. Brain Research, 405(2), 253–267. https://doi.org/10.1016/0006-8993(87)90295-2

Nobre, A. C., & Ede, F. van. (2018). Anticipated moments: Temporal structure in attention. Nature Reviews Neuroscience, 19(1), 34–48. https://doi.org/10.1038/nrn.2017.141

Oostenveld, R., & Praamstra, P. (2001). The five percent electrode system for high-resolution EEG and ERP measurements. Clinical Neurophysiology, 112(4), 713–719. https://doi.org/10.1016/S1388-2457(00)00527-7

Oosterhof, N. N., Connolly, A. C., & Haxby, J. V. (2016). CoSMoMVPA: Multi-Modal Multivariate Pattern Analysis of Neuroimaging Data in Matlab/GNU Octave. Frontiers in Neuroinformatics, 10. https://doi.org/10.3389/fninf.2016.00027

Robinson, A. K., Grootswagers, T., & Carlson, T. A. (2019). The influence of image masking on object representations during rapid serial visual presentation. NeuroImage, 197, 224–231. https://doi.org/10.1016/j.neuroimage.2019.04.050

Robinson, A. K., Grootswagers, T., Shatek, S. M., Gerboni, J., Holcombe, A. O., & Carlson, T. A. (2020). Now you see it, now you don’t: Overlapping neural representations for the position of visible and invisible objects. BioRxiv, 2020.03.02.974162. https://doi.org/10.1101/2020.03.02.974162

Rohenkohl, G., Cravo, A. M., Wyart, V., & Nobre, A. C. (2012). Temporal Expectation Improves the Quality of Sensory Information. Journal of Neuroscience, 32(24), 8424–8428. https://doi.org/10.1523/JNEUROSCI.0804-12.2012

Rouder, J. N., Speckman, P. L., Sun, D., Morey, R. D., & Iverson, G. (2009). Bayesian t tests for accepting and rejecting the null hypothesis. Psychonomic Bulletin & Review, 16(2), 225–237. https://doi.org/10.3758/PBR.16.2.225

Rugg, M. D., Milner, A. D., Lines, C. R., & Phalp, R. (1987). Modulation of visual event-related potentials by spatial and non-spatial visual selective attention. Neuropsychologia, 25(1), 85–96. https://doi.org/10.1016/0028-3932(87)90045-5

Rungratsameetaweemana, N., Itthipuripat, S., Salazar, A., & Serences, J. T. (2018). Expectations Do Not Alter Early Sensory Processing during Perceptual Decision-Making. Journal of Neuroscience, 38(24), 5632–5648. https://doi.org/10.1523/JNEUROSCI.3638-17.2018

Smout, C. A., Tang, M. F., Garrido, M. I., & Mattingley, J. B. (2019). Attention promotes the neural encoding of prediction errors. PLOS Biology, 17(2), e2006812. https://doi.org/10.1371/journal.pbio.2006812

Wetzels, R., Matzke, D., Lee, M. D., Rouder, J. N., Iverson, G. J., & Wagenmakers, E.-J. (2011). Statistical Evidence in Experimental Psychology: An Empirical Comparison Using 855 t Tests. Perspectives on Psychological Science, 6(3), 291–298. https://doi.org/10.1177/1745691611406923

Zhang, W., & Luck, S. J. (2009). Feature-based attention modulates feedforward visual processing. Nature Neuroscience, 12(1), 24–25. https://doi.org/10.1038/nn.2223

Zuanazzi, A., & Noppeney, U. (2018). Additive and interactive effects of spatial attention and expectation on perceptual decisions. Scientific Reports, 8. https://doi.org/10.1038/s41598-018-24703-6

